# Structural basis for the differential regulatory roles of the PDZ domain in C-terminal processing proteases

**DOI:** 10.1101/621037

**Authors:** Chuang-Kai Chueh, Nilanjan Som, Lu-Chu Ke, Meng-Ru Ho, Manjula Reddy, Chung-I Chang

## Abstract

Carboxyl (C)-terminal processing proteases (CTPs) participate in protective and regulatory proteolysis in bacteria. The PDZ domain is central to the activity of CTPs but plays inherently different regulatory roles. For example, the PDZ domain inhibits the activity of the signaling protease CtpB by blocking the active site but is required for the activation of Prc (or Tsp), a tail-specific protease that degrades the ssrA-tagged proteins. Here, by structural and functional analysis we show that in the unliganded resting state of Prc, the PDZ domain is docked inside the bowl-shaped scaffold without contacting the active site, which is kept in a default misaligned conformation. In Prc, a hydrophobic substrate sensor distinct from CtpB engages substrate binding to the PDZ domain and triggers a structural remodeling to align the active site residues. Therefore, this work reveals the structural basis for understanding the contrasting roles of the PDZ domain in the regulation of CTPs.

## Introduction

In Gram-negative bacteria, the periplasm is a multifunctional compartment between the porous outer membrane and the inner (or cytoplasmic) membrane. Similar to the eukaryotic endoplasmic reticulum, the bacterial periplasm provides essential functions including transport, folding, oxidation, and quality control of proteins and lipoproteins; it also confers mechanical strength to the cell by synthesizing the polymeric peptidoglycan (PG), an important structural element of the cell wall (Miller and Salama, 2018).

Prc (also named Tsp) is a member of the family of C-terminal processing proteases (CTPs) (Rawlings et al., 2002), which is featured by an embedded regulatory PDZ-domain inserted into a serine protease domain (Hara et al., 1991; Su et al., 2017). CTPs are located in the periplasm of Gram-negative bacteria. In *E. coli*, Prc is involved in degrading abnormal proteins that are unfolded or partially folded and marked cotranslationally at the C-terminus by SsrA tag (Keiler et al., 1996), which are exported to the periplasm through the Sec translocon (Pugsley, 1993). Prc is also responsible for C-terminal processing of the lipoprotein penicillin-binding protein 3 (FtsI) (Hara et al., 1991), a key component of the divisome that catalyzes the cross-linking of PG during cell division (Karimova et al., 2005; Sauvage et al., 2014). By associating with its adaptor protein, NlpI, Prc is also involved in the regulated proteolysis of MepS, a lipid-anchored hydrolase specific for PG cross-links (Singh et al., 2015). The protease activity of Prc in the periplasm contributes to bacterial evasion of killing by the host complement system (Wang et al., 2012). Deletion of *prc* results in altered cell morphology, temperature-sensitive growth under osmotic stress, reduced heat-shock response, leakage of periplasmic proteins, increased antibiotic sensitivity, and reduced virulence (Deng et al., 2018; Hara et al., 1991; Liao et al., 2016; Seoane et al., 1992; Wang et al., 2012), which are in accordance with the role of Prc in processing the periplasmic lipoproteins involved in PG synthesis.

The PDZ domain of CTPs is responsible for substrate binding and involved in regulating the activity of CTPs. In the signaling protease CtpB, which forms an intertwined dimeric ring, the PDZ domain plays an inhibitory role by physically blocking the proteolytic active site, thereby disrupting the catalytic triad; deletion of the PDZ domain yields a constitutively active protease (Mastny et al., 2013). Substrate binding activates CtpB by inducing the repositioning of the PDZ domain away from the proteolytic site, which is mediated by a polar amino acid from the PDZ domain as the substrate sensor (Mastny et al., 2013).

By contrast, Prc forms a monomeric bowl-like structure with an attached PDZ domain (Su et al., 2017). The activity of Prc strictly requires the PDZ domain and the PDZ deletion abolishes completely the protease activity (Su et al., 2017). Moreover, a pair of hydrophobic residues located in the hinged region connecting to the PDZ domain is proposed to be the substrate sensor mediating the activation of Prc by substrate binding (Su et al., 2017). Therefore, the substrate-triggered activation of Prc is likely to be mediated by the PDZ domain through a structural mechanism distinct from that of CtpB.

To understand the structural basis for the activating role of the PDZ domain in regulating the activity of Prc, we have determined a set of crystal structures in the unliganded resting-state of Prc, alone or in complex with NlpI, with deleted PDZ domain or side-directed mutations in either the PDZ ligand-binding site or the substrate-sensing hinge. In the unliganded resting state, the lid-like PDZ domain is positioned inside the bowl-like body but does not make contact with the proteolytic site. In the structure, the hinge region is deformed into coils and the proteolytic active site residues are misaligned. Similar inactive conformation of the proteolytic site is also seen in the structures of Prc with the PDZ deletion or mutated hinge residues. Structural comparison to the substrate-bound activated structure of Prc reveals how substrate binding to the PDZ domain induces extensive alignment of the proteolytic active site residues mediated by structural rearrangement of the substrate-sensing hinge. Overall, these results provide the structural basis for understanding the contrasting regulatory roles of the PDZ domain in the activation of different CTPs.

## Results

### Characterizing Prc with mutations in the substrate binding sites

In order to capture Prc in the unliganded resting state by preventing substrate binding to the PDZ domain and the proteolytic site, we have engineered five Prc mutants as follows: (1) Prc-ΔPDZ (residues 246-339 deleted)(Su et al., 2017), (2) Prc-L252Y with the mutation designed to block the PDZ ligand-binding pocket, (3) Prc-K477A/L252Y with an addition mutation on the catalytic residue Lys477, (4) Prc-S452I/L252Y, where the catalytic residue Ser452 is mutated to isoleucine with a bulky hydrophobic side chain, and (5) Prc-L245A/L340G with double mutations on the critical substrate-sensing hinge (Su et al., 2017). We next compared the proteolytic activity of these Prc mutants with wild-type (WT) Prc against two *in vivo* substrates, FtsI and MepS (Hara et al., 1991; Singh et al., 2015). Using purified recombinant FtsI, we showed that it is cleaved by WT Prc into a shorter processed form, supporting previous finding (Hara et al., 1991)(Fig. 1A). Interestingly, Prc could not cleave FtsI in the presence of NlpI, which is needed for efficient degradation of MepS (Fig. 1A)(Singh et al., 2015; Su et al., 2017); presumably, the three-sided MepS-docking cradle formed by bound NlpI excludes the access of the larger sized FtsI to Prc. By contrast, Prc-ΔPDZ, Prc-L252Y, and Prc-L245A/L340G all failed to process FtsI (Figs. 1B and C). All double Prc mutants also lost MepS degrading activity (Fig. 1D). Viability assays also showed that all the double-mutation alleles did not complement a *prc* deletion mutant (Fig. 1E). Finally, analytical ultracentrifugation (AUC) analysis indicated a smaller particle size of the double mutants compared to Prc-K477A, which is locked in the substrate-bound activated open conformation (Su et al., 2017)(Fig. 1F), suggesting that these double mutants may adopt a more compact structure than Prc-K477A. Indeed, thermal shift assays showed single melting transitions for these double mutants distinct from the two-phased Prc-K477A (Fig. 1G).

**Figure 1.**
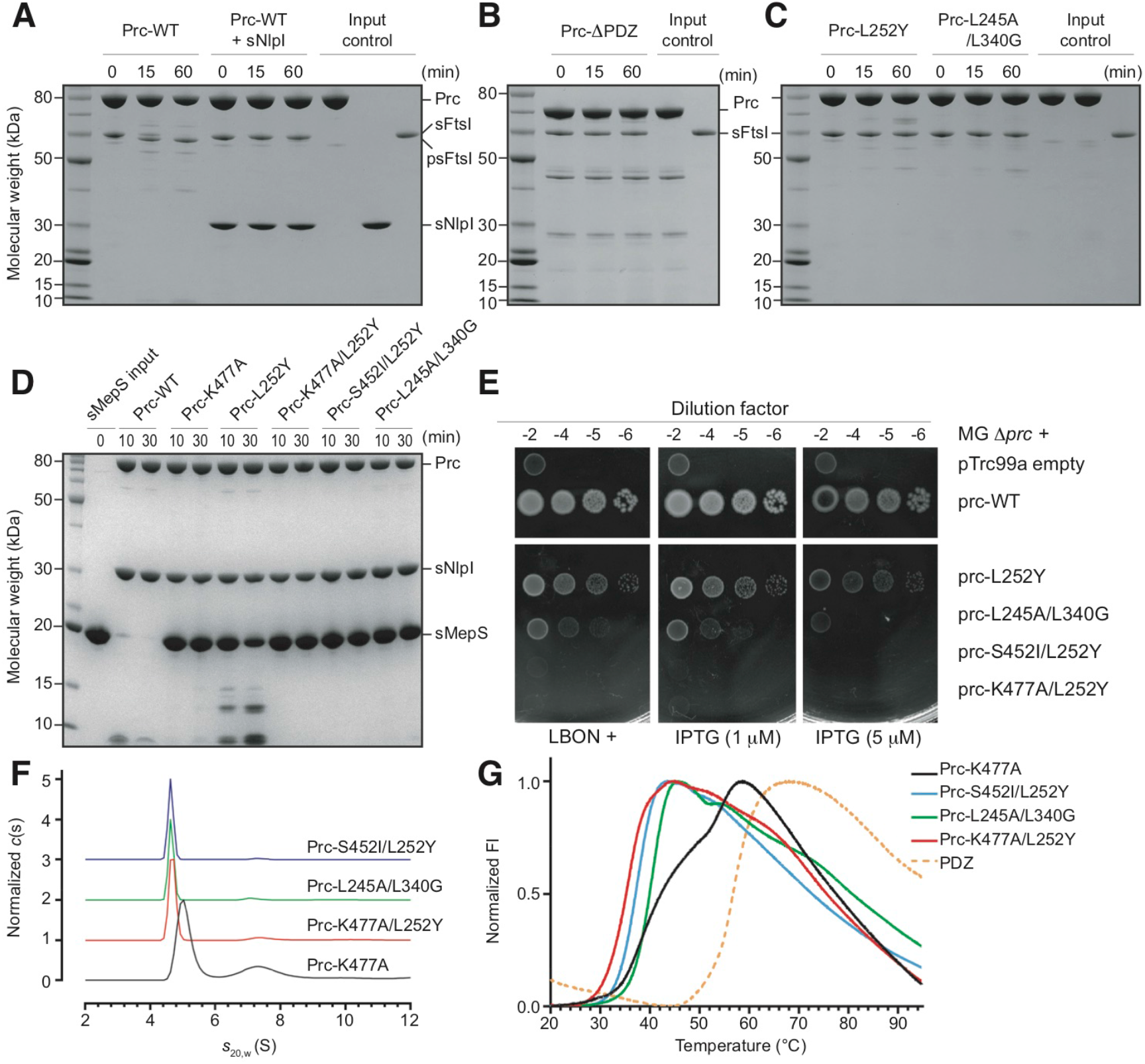
Biochemical and biophysical properties of Prc and various mutants. **(A)** SDS-PAGE analysis monitoring the cleavage of FtsI (residues 57-588; sFtsI) without the N-terminal transmembrane segment into the processed form (residues 57-577; psFtsI) by wild-type Prc, with and without NlpI. **(B)** Cleavage assays by Prc with deleted PDZ domain. **(C)** Cleavage assays by Prc with mutations in the PDZ ligand-binding pocket (L252Y) and the substrate-sensing residues (L245A and L340G). **(D)** Degradation assays of MepS by wild-type Prc and various Prc mutants. **(E)** Viability assays examining the effects of various Prc mutations on the growth of Δ*prc* mutant on media of low osmotic strength (LBON). The wild type *prc* in pTRC99a plasmid has a leaky expression sufficient to complement Δ*prc* mutant even without the inducer. **(F)** Sedimentation velocity profiles comparing the molecular sizes of various Prc double mutants and Prc-K477A, which is in the liganded activated state (Su et al., 2017). **(G)** DSF melting curves comparing the melting transitions of the Prc double mutants with liganded Prc-K477A and the isolated PDZ domain.

### Structure of Prc-S452I/L252Y in the unliganded resting state

According to the above results, we performed crystallization screening experiments for each of the Prc double mutants that likely adopt unliganded resting-state conformation, alone or in complex with NlpI. We determined the structure of Prc-S452I/L252Y (Table S1), which is trapped in the unliganded resting state. Prc has a platform-like protease domain harboring the proteolytic groove, which in the activated state is enclosed by a vault-like structural element comprising helix h9 and a three-stranded b2-b19-b20 antiparallel β-sheet (Su et al., 2017). In addition to the conserved protease domain, Prc contains extended N-terminal and C-terminal helical domains (named NHD and CHD, respectively), which are adjoined together via two β-strands (Fig. 2A). In Prc, the vaulted protease domain, NHD, and CHD form a round bowl-like structure. The PDZ domain is inserted into the protease domain between helix h9 and the platform (Figs. 2A and 2B). In the structure of Prc-S452I/L252Y, the PDZ domain is docked inside the bowl (Fig. 2A), interacting with residues dispersed across a wide region mainly from NHD and CHD (Fig. S1A); these residues constitute a total interface area of 1595 Å^2^. Almost half (49.4%) of the surface residues of the PDZ domain interacting with the bowl are distributed evenly across the surface (Fig. S1B).

**Figure 2.**
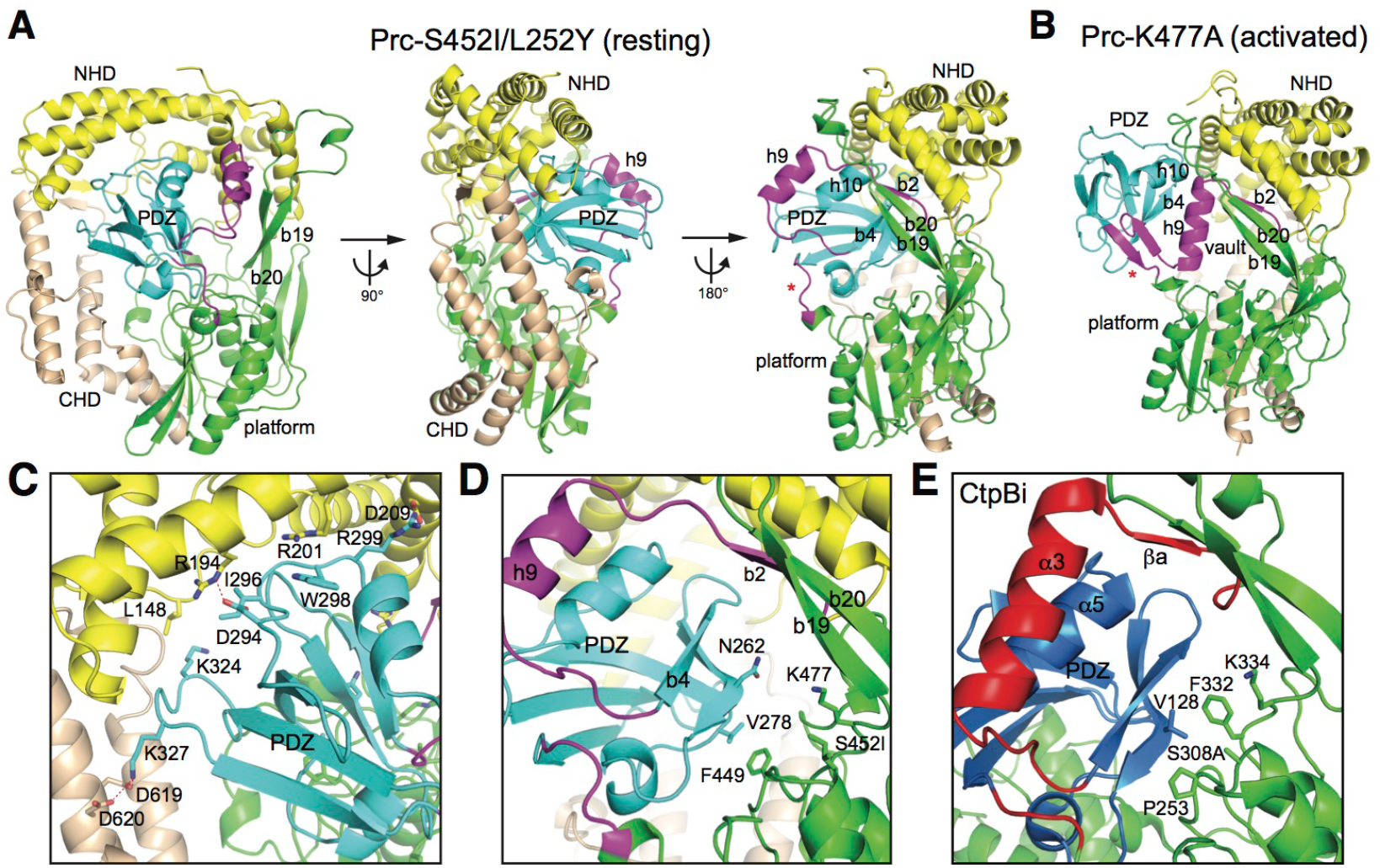
Overall structure of Prc in the resting state. **(A)** Three orthogonal views of Prc-S452I/L252Y in ribbon representations. **(B)** For comparison, the structure of Prc-K477A (bound substrates were omitted) determined previously in the activated state was shown in the same orientation as the right view in (A). **(C-D)** Zoom-in views showing the interaction of the PDZ domain with NHD and CHD (C) and with the protease platform (D). **(E)** Zoom-in view showing the PDZ domain interaction in the inhibited resting CtpB (CtpBi). The associated N- and C-terminal helical domains (NHD and CHD), the PDZ domain, and the protease platform domain are indicated and shown in different colors. The vault element consisting of helix h9 and strand b2 and the hinge coil (indicated by an asterisk) undergoing remodeling during ligand-dependent activation are highlighted in magenta.

### Intramolecular interaction of the docked PDZ domain in Prc

In the resting structure, the PDZ domain is surrounded by NHD, CHD, and the protease domain. Most of the contacting residues of the PDZ domain are from the loops and are polar amino acids. The interacting residues of NHD and CHD are mostly polar amino acids from the helices and several hydrophobic residues from the loops. However, few are involved in specific side-chain interactions except for the polar pairs of Lys327-Asp619, Arg299-Asp209, and Arg194-Asp294 (Fig. 2C). Instead, many residues engage side-chain stacking interaction to form Van der Waals contacts. Perhaps owing to the non-specific nature of the docking interaction, the PDZ domain in the resting structure shows higher temperature factor (B-factor) values than the bowl-shape scaffold of Prc.

Interestingly, the docked PDZ domain makes little contact with the protease domain; the only contact is made between the stacking side chains of PDZ-Val278 and Phe449 at the back, which anchors the PDZ domain in a specific position to expose the ligand-binding strand b4 (Fig. 2D). However in CtpB, helix α5 of the inhibitory PDZ domain packs against helix α3 and strand βa. Importantly, the PDZ residue Val128 blocks the catalytic Ser308 by binding to a hydrophobic pocket walled by Pro253 and Phe332 (Fig. 2E), but the corresponding PDZ residue in Prc, Asn262, makes no contact whatsoever (Fig. 2D). Therefore, the lack of extensive interaction of the PDZ domain with the protease domain and of any direct contact of the PDZ domain with the proteolytic active site in the resting structure of Prc is significantly different from that of CtpB, which further supports that the PDZ domain of Prc does not assume an inhibitory function (Su et al., 2017).

### Structural difference between liganded activated and unliganded resting Prc

Comparing to the liganded activated state, in the unliganded resting structure the docked PDZ domain is repositioned to expose its ligand-binding site for the substrate C-terminus to an open space above the platform (Fig. 3A). The two-β-stranded substrate-binding hinge, formed in the activated state and connecting the PDZ domain to helix h9 and the platform, is unfolded into two coils; the critical substrate sensing residues Leu245 and Leu340 are separated and become solvent-exposed (Fig. 3B). Helix h9 is drifted away and partially unfolded to make the large vaulted space above the platform (Fig. 3B). Lastly, without a folded connecting hinge the proteolytic platform is angled down with misaligned active site residues: the catalytic Lys477 and Ser452 are apart from each other, and the amide groups of Ala453 and Gly398 forming the oxyanion pocket are out of place (Figs. 3B and C).

**Figure 3.**
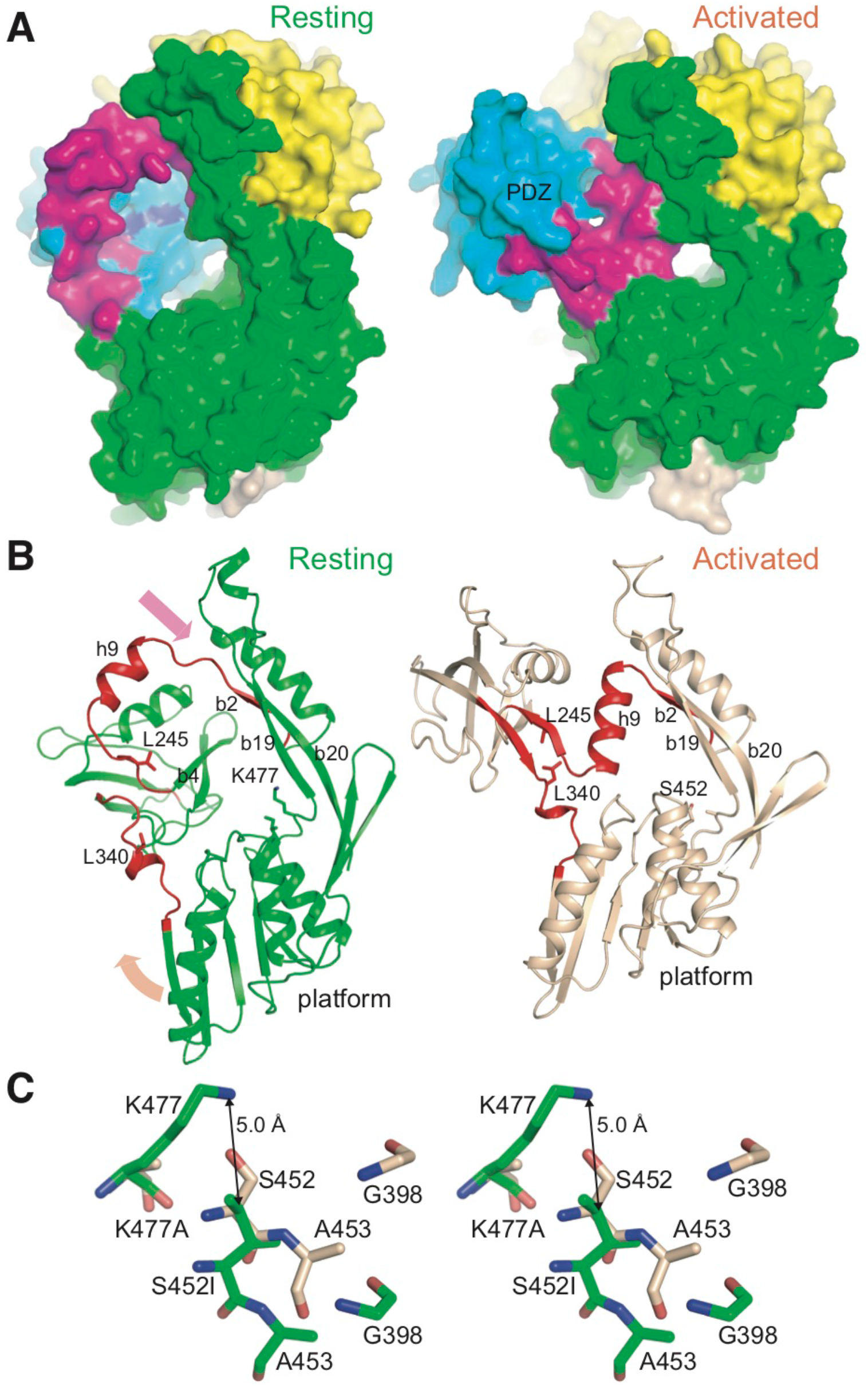
Structural difference between the resting and activated states of Prc. **(A)** Structures of Prc-S452I/L252Y (left) and Prc-K477A (right; bound substrate omitted) are shown in surface representations in similar orientation. The coloring scheme is the same as in Fig. 2. The PDZ ligand-binding site is highlighted by coloring L252Y in blue. **(B)** Structural comparison of the resting and activated Prc highlighting the remodeling the two hinge coils (colored in red) into a pair of β-strands during activation; the arrows indicate the directions of movement of helix h9 and the protease platform. The NHD and CHD were removed for clarity. **(C)** Stereo view of the catalytic residues Lys477 and Ser452, and the oxyanion hole residues Gly398 and Ala453 in the two states. The superposition was obtained by structural alignment of Prc-S452I/L252Y (resting form; green color; chain A) and Prc-K477A (activated form; wheat color; PDB code 5WQL_C)(2.14 Å rmsd; 503 residues aligned).

### Structures of Prc without the PDZ domain or the hydrophobic sensor

To assess the structural role of the PDZ domain and the hydrophobic sensor residues, which have been shown to be required for Prc activity, we also determined the crystal structures of Prc-ΔPDZ and Prc-L245A/L340G, both are in complex with NlpI. In both structures, helix h9 and the hinge region are disordered and invisible from the electron-density map. Additionally, the entire PDZ domain in the structure of Prc-L245A/L340G is also missing in the electron-density map. Nevertheless, the structures of the bowl-like body of Prc-ΔPDZ and Prc-L245A/L340G including the protease platform are superimposable to that of the resting Prc-S452I/L252Y (Fig. S2A and Table S2). The proteolytic active site residues in Prc-ΔPDZ and Prc-L245A/L340G are in the misaligned resting-state conformation (Fig. S2B). These structures confirm that, without the PDZ domain or the hydrophobic substrate sensor, Prc is maintained in the inactive resting state.

### Flexibility of helix h9 in the resting state

We also crystallized Prc-S452I/L252Y bound to NlpI in the space group of P2_1_2_1_2_1_, which differs from that of Prc-S452I/L252Y alone (P3_2_21) but is the same as that of the substrate-bound NlpI-Prc-K477A crystals trapped in the activated state (Su et al., 2017). The crystal contacts of these P2_1_2_1_2_1_ forms involve the NlpI dimer and the NHD and CHD of Prc only; hence, they provide further information about the structural difference between the two conformational states. The structure of Prc-S452I/L252Y in NlpI-bound complex shows a similar resting-state conformation, with the RMSD value of 0.62 comparing to that of Prc-S452I/L252Y alone (Table S2). Interestingly, unlike the liganded Prc (Fig. 4A), the NlpI complexed Prc-S452I/L252Y shows flexible helix h9 with poor electron-density map and the coiled region connecting to the PDZ domain are partially disordered (Fig. 4A). Moreover, the docked PDZ domain in these resting structures has higher B-factor values than the protease domain, which is in sharp contrast to the liganded activated structure showing relatively rigid PDZ domain, helix h9, and the substrate-sensing hinge (Figs. 4A and B). To probe the flexibility of h9 in these resting-state mutants in solution, we performed limited proteolysis using V8 protease, which has been shown to cleave Prc at the peptide bond between Asn211 and Thr212 of helix 9, yielding two fragments of molecular masses of 49.5 and 25.7 kDa (Beebe et al., 2000). We found that limited V8 proteolysis indeed resulted in the two fragments from Prc-S452I/L252Y but not in Prc-K477A at various incubation times (Fig. 4C), supporting the flexibility of helix h9 in the resting state.

**Figure 4.**
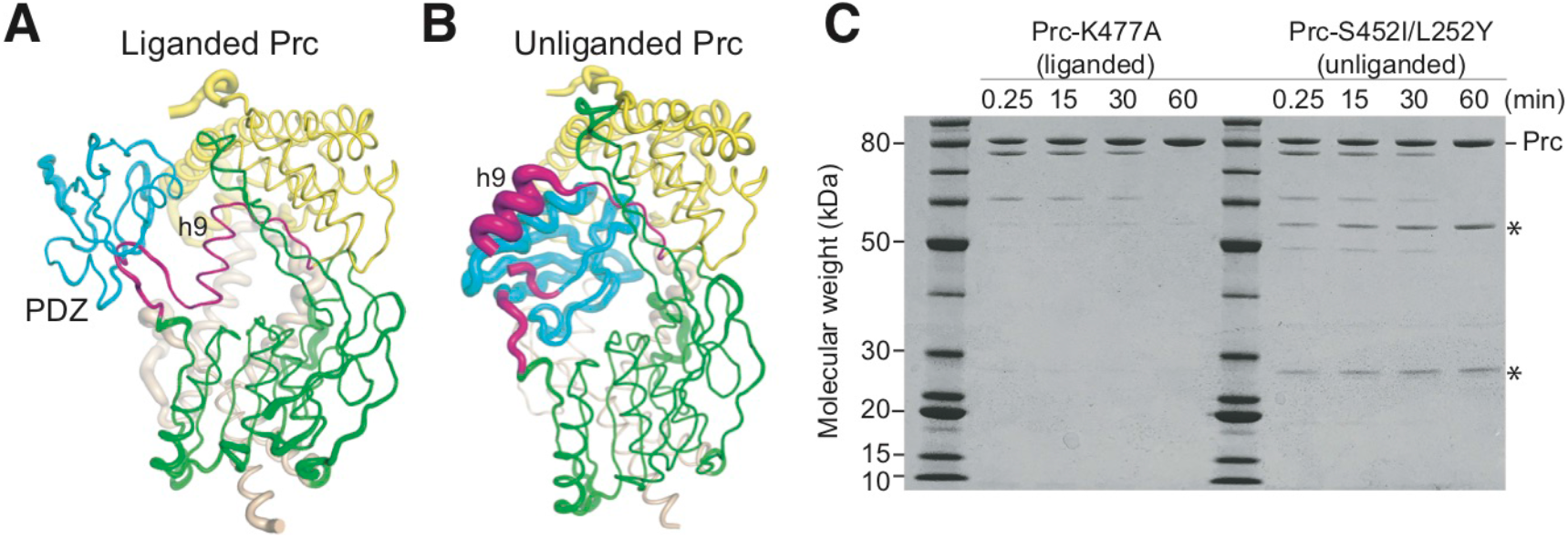
Flexibility of helix h9 in the resting state. **(A-B)** Structures of NlpI-bound NlpI-Prc-K477A (A) and NlpI-Prc-S452I/L252Y (B) in tube representations with the tube diameter correlated to the b-factor of the structure. The NlpI dimer was omitted for clarity. The coloring scheme is the same as in Fig. 2. **(C)** SDS-PAGE analysis monitoring the cleavage of Prc by V8 protease at the indicated time points. The two fragments generated by cleavage at residue Asn211 of helix h9 are indicated by asterisks.

## Discussion

The work presented here has revealed the structural basis for the activating role of the PDZ domain in Prc. Our results show that in unliganded resting state, Prc forms an inactive structure characterized by a deformed proteolytic groove resulting from misalignment of the loops forming the catalytic Lys-Ser dyad and the oxyanion pocket. In the resting state, the PDZ domain of Prc is docked inside a bowl-like scaffold with the ligand-binding site exposed. Notably, the PDZ domain engages intramolecular interaction mainly with the NHD and CHD domains rather than with the catalytic active site in the protease domain. In the absence of the PDZ domain, Prc-ΔPDZ also adopt the resting-state structure with a similarly misaligned proteolytic domain. These results support the finding that the PDZ domain regulates the protease activity of Prc by serving as an activator but not an inhibitor. Prc activation is achieved by substrate binding to the PDZ domain, which is sensed by conserved hydrophobic residues Leu340 and Leu254 to stabilize the substrate-bound active conformation. The repositioned PDZ domain induces extensive remodeling of the functional proteolytic platform to enable substrate cleavage reaction (Fig. 5A).

**Figure 5.**
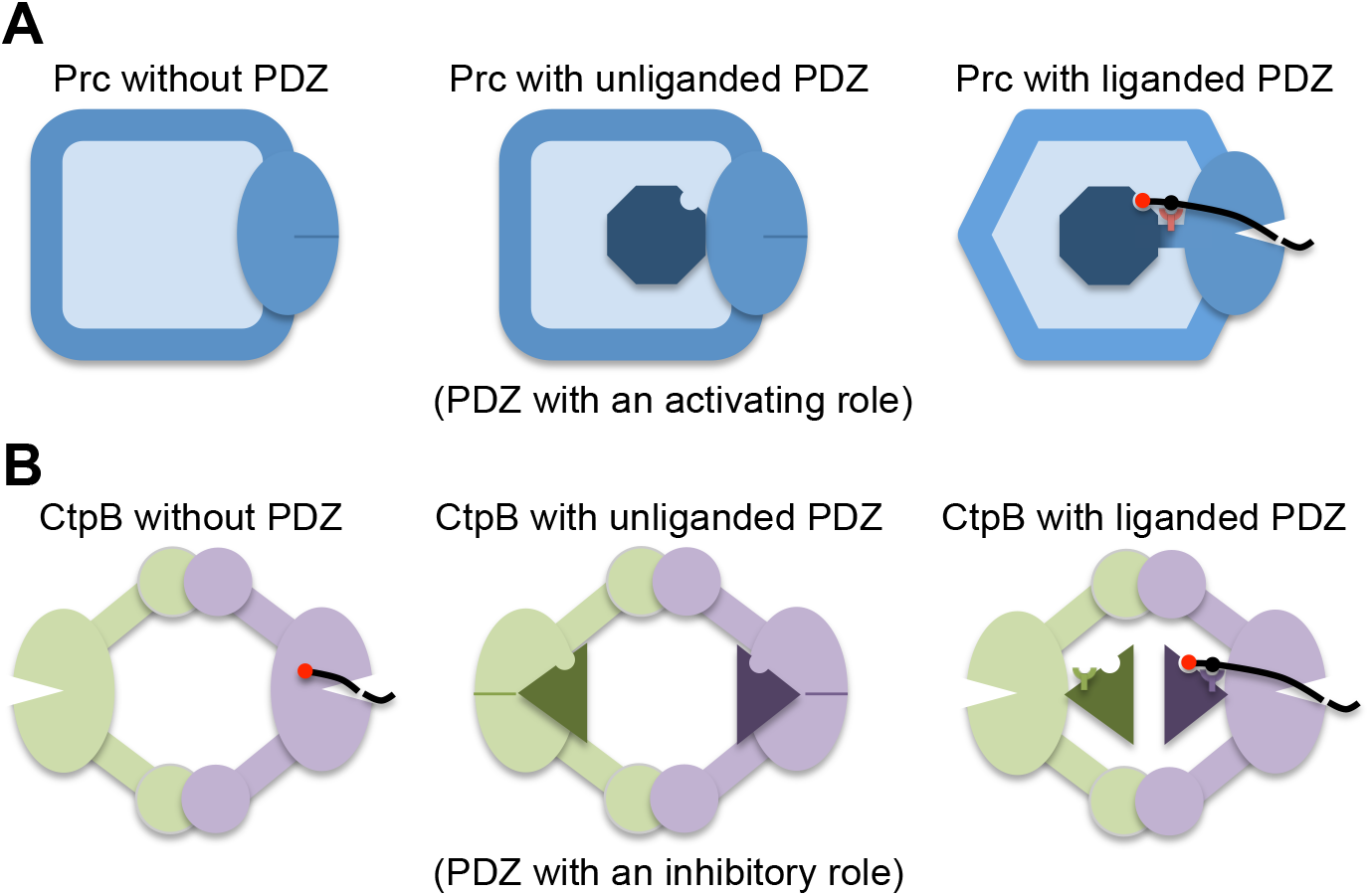
Comparison of the structural mechanisms for the regulation of the protease activity of Prc and CtpB by the PDZ domain. **(A)** In Prc, the PDZ deletion results in an inactive protease (left), as evidenced by a misaligned protease active site (indicated by a solid dash). In the resting state, unliganded PDZ domain (the octagon) docks inside the bowl-like scaffold structure of Prc and makes no contact with the protease active site (middle). Upon substrate binding, the hydrophobic sensor (Leu340; indicated by a Y-shaped symbol) engages the bound substrate and triggers structural remodeling to align the protease active site for substrate cleavage (right). **(B)** In dimeric CtpB, deletion of the PDZ domain yields a constitutively active protease (left). In the inhibited resting state, the protease active site is disrupted by the docked PDZ domain (the triangles)(middle). Substrate binding induces repositioning of the PDZ domain, stabilized by the polar sensor (Arg168) from the PDZ domain (right).

The mechanism of protease activation by the PDZ domain in Prc shown here is fundamentally different from that in CtpB, reported previously (Mastny et al., 2013). In CtpB, the protease is active without the PDZ domain. In the resting state, the PDZ domain of CtpB binds to the protease domain and physically disrupts the catalytic active site. Binding of the substrate to the PDZ domain is sensed by PDZ residue Arg168, which stabilizes the reposition of the PDZ domain away from the protease domain to relieve the inhibition effect (Fig. 5B).

The different roles of the PDZ domain in the activation of Prc and CtpB may be paralleled with that for the structurally unrelated HtrA-family proteases DegP and DegS, in which oligomerization involving the PDZ domains contributes additional regulatory roles (Clausen et al., 2011). DegP requires the PDZ domain for activation (Iwanczyk et al., 2007), but the PDZ domain in DegS mainly inhibits the activity (Sohn et al., 2007; Walsh et al., 2003). Therefore, the PDZ domain can inherently bring different effects in the activity regulation of the PDZ-containing proteases.

Many of the structural features of the unliganded Prc and conformational changes induced by substrate C-terminus are also different from CtpB. Helix α3 in CtpB, equivalent to helix h9 of Prc, is structured in both resting and activated states. In the two states of CtpB, the PDZ domain maintains critical contact with helix α3 by the sensor Arg168 (Mastny et al., 2013). By contrast, the PDZ domain of Prc does not interact with helix h9 in the resting state. Notably, as helix α3 is not partially unfolded in resting CtpB, the open gate framed by the helix in CtpB is not as large as that in the unliganded Prc (Figs. S3A and S3B). Interestingly, a structure of CtpA determined in the inactive state shows an even smaller gate (Figs. S3C and S3D)(Liao et al., 2000). Therefore, the large open gate in Prc, framed by the disordered helix h9, may allow access of folded or unfolded substrates with varied lengths of unstructured C-terminal tails. By contrast, CtpA and CtpB are known to recognize and process specific substrates: SpoIVFA and photosystem II D1 protein, respectively, explaining their smaller open gates in the resting state. Lastly, CtpB forms a pseudo-symmetric ring-like homodimer locked via their N- and C-terminal dimerization domains, which may permit only limited conformational change triggered by the liganded PDZ domain if only one of its two interlocked subunits are activated by a substrate. However, given the primary inhibitory role of the PDZ domain in CtpB, a substrate-triggered release of the PDZ domain from one catalytic active site may be sufficient to active CtpB without the need of larger conformational change of the PDZ and the protease domains shown in the activation of Prc (Su et al., 2017).

Our crystallographic results also reveal that Prc-L245A/L340G, without the two ligand-sensing residues, has a flexible PDZ domain missing in the electron density map; importantly, the bowl adopts a resting-state structure showing a dis-localized protease platform and deformed active site loops, which are superimposable to the structure of unliganded Prc-S452I/L252Y (Fig. S2A). Since this mutant contains normal PDZ ligand-binding and catalytic residues but has lost completely the ability to degrade MepS or process FtsI, its structure demonstrates that the two hydrophobic residues are important substrate sensors essential for stabilizing the ligand-bound activated conformation of Prc.

Unexpectedly, our structural and limited proteolysis results show that helix h9 of Prc is flexible in the resting state but becomes structurally defined to narrowly enclose the bound cleavage-site substrate polypeptide in the activated state. The disorder-to-order transition of helix h9, which completes the active site, regulated allosterically by liganded PDZ domain, is therefore different from helix α3 of CtpB but similar to that of the active site loops of the structurally unrelated high temperature requirement A (HtrA) family of oligomeric PDZ-proteases, such as DegP and DegS (Clausen et al., 2011). However, the active site of HtrA proteases is exposed and does not enclose the substrate; interestingly, the PDZ activating ligand and the cleavage-site substrate bound in the same subunit come from different polypeptide chains and are thus not covalently linked (Kim et al., 2011; Sohn et al., 2007). By contrast, binding of the C-terminus of a substrate to the PDZ domain through the large open gate of Prc must always result in the entrapment of the same substrate polypeptide at the active site enclosed by helix h9. Therefore, the activation mechanism of Prc combines features from those of CtpB and HtrA proteases.

Finally, our structural studies have provided mechanistic insight into the operation of Prc by the activating PDZ domain undergoing relocation upon substrate binding (Movie S1). During activation, helix h9 moves in a direction opposite to that of the liganded PDZ domain and assumes a center position between the two substrate-binding sites, in the PDZ domain and the proteolytic groove (Figs. S4A and B). Concurrently, the flexible helix h9 undergoes its own remodeling and a disorder-to-order transition, which serve to completely enclose the substrate polypeptide bound to the proteolytic groove (Fig. 4C). As such, helix h9 may have a pulley-like function to convert the conformational change of the substrate-bound PDZ domain into driving substrate translocation for Prc to degrade folded or incompletely folded protein substrates *in cis*.

## Materials and Methods

### Construction of BL21 (λDE3) *prc*::Cm mutant

P1-phage mediated transduction was performed as described (Miller, 1992). P1 phage lysate was generated using MG1655 *prc*::Tn*10*dCm strain (from our laboratory collection) and the *prc*::Tn*10*dCm mutation was transferred by P1 transduction into BL21 (λDE3) and selected on Chloramphenicol containing LB plates (15 µg/mL). This mutant behaves identical to that of a *prc* deletion mutant.

### Cloning and mutagenesis

The Prc mutants (Prc-K477A, Prc-L252Y, Prc-S452I, Prc-K477A/L252Y, Prc-S452I/L252Y, Prc-L245A/L340G) were generated by PCR-based site-directed mutagenesis (primers listed in Table S3), using wild-type Prc plasmid as the template (Su et al., 2017). Prc-ΔPDZ (Δ247-339) with a C-terminal His-tag was also cloned into pTrc99A vector. As for sNlpI and sMeps, the encoding DNA sequences without lipoprotein signal peptides were cloned into pET28a and pET21a vector, respectively. sMepS was cloned with a C-terminal His-tag, while sNlpI was expressed with a N-terminal His-tag and tobacco etch virus (TEV) protease cleavage site. All of the constructs were sequenced before follow-up experiments. MG1655 genomic DNA was used as a template to PCR amplify the *ftsI* gene encoding residues 57-588 without the N-terminal transmembrane helix (sFtsI) (Table S3). The amplified product and pET28a vector were digested with NheI and BamHI restriction enzymes (New England Biolabs, USA), purified, and ligated using T4 DNA ligase (New England Biolabs, USA). The positive clones were identified by colony PCR and confirmed by sequence analysis.

### Protein expression and purification

To prevent contamination or pre-processing by endogenous Prc, Prc mutant proteins and sFtsI were expressed in *E. coli* ΔPrc cells (strain MR812 and BL21 (λDE3) *prc*::Cm, respectively)(Su et al., 2017). Full-length Prc, sNlpI and sMepS were expressed in *E. coli* BL21 (λDE3) cells as described previously (Su et al., 2017). Cells were grown in LB medium until the optical density at 600 nm reached 0.6 - 0.8, and were induced with 1 mM isopropyl β-D-thiogalactopyranoside (IPTG) for 4 hrs at 22°C. Cell pellets were collected after centrifugation and resuspended in lysis buffer containing 50 mM Tris-HCl (pH 8.0) and 500 mM NaCl. After being ruptured by French press (Avestin) and centrifuged at 35,000 x g, the supernatants were collected and incubated with Ni-NTA resins (QIAGEN) for 2 hrs at 4°C. Proteins were further washed with a stepwise imidazole gradient and eluted with 250 mM imidazole.

All of the recombinant proteins were dialyzed to remove imidazole against different buffer components. For further assays, Prc and sNlpI were dialyzed against buffer containing 20 mM Tris-HCl (pH 8.0) and 150 mM NaCl. sMepS was dialyzed against same buffer components with additional 2 mM dithiothreitol (DTT). sFtsI, on the other hand, was dialyzed against buffer containing 20 mM HEPES (pH 7.0) and 150 mM NaCl. Prc and sNlpI were further purified by MonoQ 5/50 GL column chromatography (GE Healthcare) at pH 8.0. For protein crystallization, most of the Prc mutants (Prc-K477A/L252Y, Prc-S452I/L252Y, Prc-PDZ) and sNlpI were first dialyzed against buffer containing 25 mM Tris-HCl (pH 8.0) and 50 mM NaCl. Prc-L245A/L340G was dialyzed against buffer containing 25 mM Tris-HCl (pH 8.0) and 150 mM NaCl after 250 mM imidazole elution. All of the recombinant proteins were then purified by MonoQ 5/50 GL column chromatography at pH 8.0. After that, Prc and sNlpI were concentrated, mixed in 1:2 molar ratio, and subjected to a Superdex 200 10/300 GL column (GE Healthcare) which was equilibrated with buffer containing 25 mM Tris-HCl (pH 8.0) and 150 mM NaCl to get Prc-NlpI complexes.

### Crystallization and Data collection

Hanging-drop vapor-diffusion method was performed for crystallization. Protein solutions with concentrations ranging from 10 to 20 mg/mL were mixed with equal volumes of reservoir solutions, and crystallization experiments for most of the protein samples were performed at 22°C except for the complex of sNlpI-Prc-ΔPDZ, which was performed at 16°C. For the sNlpI-Prc-L245A/L340G complex, crystals were grown with solutions containing 0.2 M sodium citrate tribasic dihydrate and 10-13% PEG3350. For sNlpI-Prc-S452I/L252Y complex, protein was crystallized with solution containing 0.1 M imidazole (pH 8.0), 0.2 M calcium acetate and 11% PEG8000. For Prc-S452I/L252Y, crystals were obtained in solution containing 0.2 M sodium thiocyanate (pH 6.4-6.8) and 20% PEG3350. For sNlpI-Prc-ΔPDZ complex, crystal was grown in solution containing 0.2 M ammonium sulfate, 0.1 M Tris-HCl (pH 8.5) and 25% PEG3350. Crystals were cryo-protected by transferring to their corresponding reservoir solutions supplemented with 20% glycerol or 20% ethylene glycol before data collection. Beside the dataset of sNlpI-Prc-L245A/L340G, which was collected at BL-1A of Photon Factory (Japan), datasets of other four protein complexes were collected at NSRRC (Taiwan). The diffraction data of sNlpI-Prc-S452I/L252Y and sNlpI-Prc-ΔPDZ were collected at beamline TPS 05A, whereas the data set of Prc-S452I/L252Y was collected at beamline TLS 15A1. All diffraction data were indexed, integrated, and scaled using HKL2000 (Otwinowski and Minor, 1997).

### Structure determination and refinement

Using the structures of sNlpI and separate domains of liganded Prc-K477A (PDB code 5WQL) as search models, the complex structure of sNlpI-Prc-S452I/L252Y was solved by molecular replacement with the program Phaser (Mccoy et al., 2007). Partial solutions for these PDZ-missing tetramer complexes were then obtained and subjected to rigid-body refinement. The final structures were built with MOLREP using two PDZ domains as search models and the difference Fourier maps as search space (Vagin and Teplyakov, 1997). Structures of Prc-S452I/L252Y, sNlpI-Prc-L245A/L340G and sNlpI-Prc-ΔPDZ were solved by molecular replacement with Phaser using the structure of sNlpI-Prc-S452I/L252Y as the search model. The crystals of Prc-S452I/L252Y were found to be merohedrally twinned with a twinning fraction 0.42 as determined by the Phenix tool Xtriage and subsequently checked by the CCP4 program TRUNCATE (Adams et al., 2010; Bailey, 1994). The CCP4 program DETWIN was then used to detwin data (Rees, 1980; Yeates, 1997). After automated model building using Phenix tool AutoBuild and the CCP4 program Buccaneer (Adams et al., 2010; Cowtan, 2006), all models were manually adjusted using Coot (Emsley and Cowtan, 2004), then iteratively refined with REFMAC5 in the CCP4 package (Murshudov et al., 2011). The final models were validated with the program MolProbity and PROCHECK (Chen et al., 2010; Laskowski et al., 1993). Crystallographic and refinement statistics are listed in Table S1.

### Degradation assays

For sFtsI degradation assays, each reaction mixture (10 μL) contained 2 μg WT or Prc variants and 0.5 μg of sFtsI in buffer containing 20 mM Tris-HCl (pH 8.0) and 150 mM NaCl. Additional sNlpI was added to Prc WT-sNlpI group, in a molar ratio of 1:1 with Prc. All of the reactions were incubated at 37°C and stopped at indicated time points by adding 5x SDS-PAGE loading dye. Samples were heated at 98°C and loaded onto 4-12% Bis-Tris gels (Invitrogen). Protein bands were then detected by Coomassie blue staining. For sMepS degradation assays, 7 μg of sMepS was incubated with 2 μg WT or Prc mutant proteins and 1 μg of sNlpI (at 1:1 molar ratio) in each reaction mixture (10 μL). Reactions were incubated and processed as described above except that the percentage of protein gels used was 12% Bis-Tris (Invitrogen).

### Viability assays

Overnight grown cultures were serially diluted in minimal media and 5 μL of each dilution was spotted on the indicated plates and incubated at appropriate temperature overnight. Plates contained 1.5% agar in either LB (1% tryptone, 0.5% yeast extract, 1% NaCl), or LBON (1% tryptone, 0.5% yeast extract).

### Analytical ultracentrifugation sedimentation velocity (AUC-SV)

AUC-SV analyses were performed in XL-A analytical ultracentrifuge equipped with a 4-hole An-60 Ti rotor and 12-mm double-sector charcoal-filled epon centerpieces (Beckman Coulter). Sedimentation velocity measurements using absorbance optics of reference buffer and samples in 20 mM Tris-HCl (pH 8.0), 150 mM NaCl were carried out at 45,000 rpm and 20°C. The buffer density and viscosity was calculated by Sednterp. Sedimentation coefficient (S) distributions were calculated using the c (s) method as implemented in the SEDFIT program (www.analyticalultracentrifugation.com) and converted to 20°C, water conditions. AUC-SV results were normalized and plotted using GraphPad Prism 7 (GraphPad Software, USA).

### Differential scanning fluorimetry (DSF)

Purified PDZ and Prc mutant proteins were diluted in 20 mM Tris-HCl (pH 8.0), 150 mM NaCl buffer to a final concentration of 0.2 mg/mL. 15 μL diluted proteins were mixed with 1 μL 100x Sypro Orange (Sigma-Aldrich) and loaded in LightCycler 480 Multiwell Plate 96 (Roche) for DSF assay. The DSF assay was conducted on a LightCycler 480 II Real-Time PCR (Roche) with the excitation and emission wavelengths set to 465 and 580 nm, respectively. Fluorescence in function of the temperature was recorded from 20°C to 95°C at 0.01°C/s rate. Melting curves were exported for further processing with GraphPad Prism 7 (GraphPad Software, USA).

### Limited proteolysis

For V8 proteolysis assay, 6 μg of Prc was incubated with 0.6 μg of V8 protease (Roche) in 0.1 M Tris (pH 7.4) at 25°C. Samples were incubated and reactions were stopped at different time points by adding 5x SDS-PAGE loading dye. After being heated at 98°C, samples were loaded onto 4-12% Bis-Tris gel. Protein bands were then detected and analyzed by Coomassie blue staining.

## Acknowledgements

We thank the synchrotron beamline supports from the Photon Factory, Japan, and NSRRC, Taiwan, ROC. This work was supported by funds from Council of Scientific and Industrial Research and Department of Biotechnology, Government of India (to M.R.), and Academia Sinica, Taiwan (to C.I.C.).

## Author contributions

C.I.C. conceived the study. C.K.C., N.S., L.C.K., M.R.H., M.R., and C.I.C. designed the experiments and analyzed the data. C.K.C. solved the crystal structures. C.K.C., N.S., L.C.K., and M.R.H. performed the experiments. C.K.C., M.R., and C.I.C. wrote the manuscript with inputs from all other authors.

## Supplemental Materials

### Supplemental Figures

**Figure S1.**
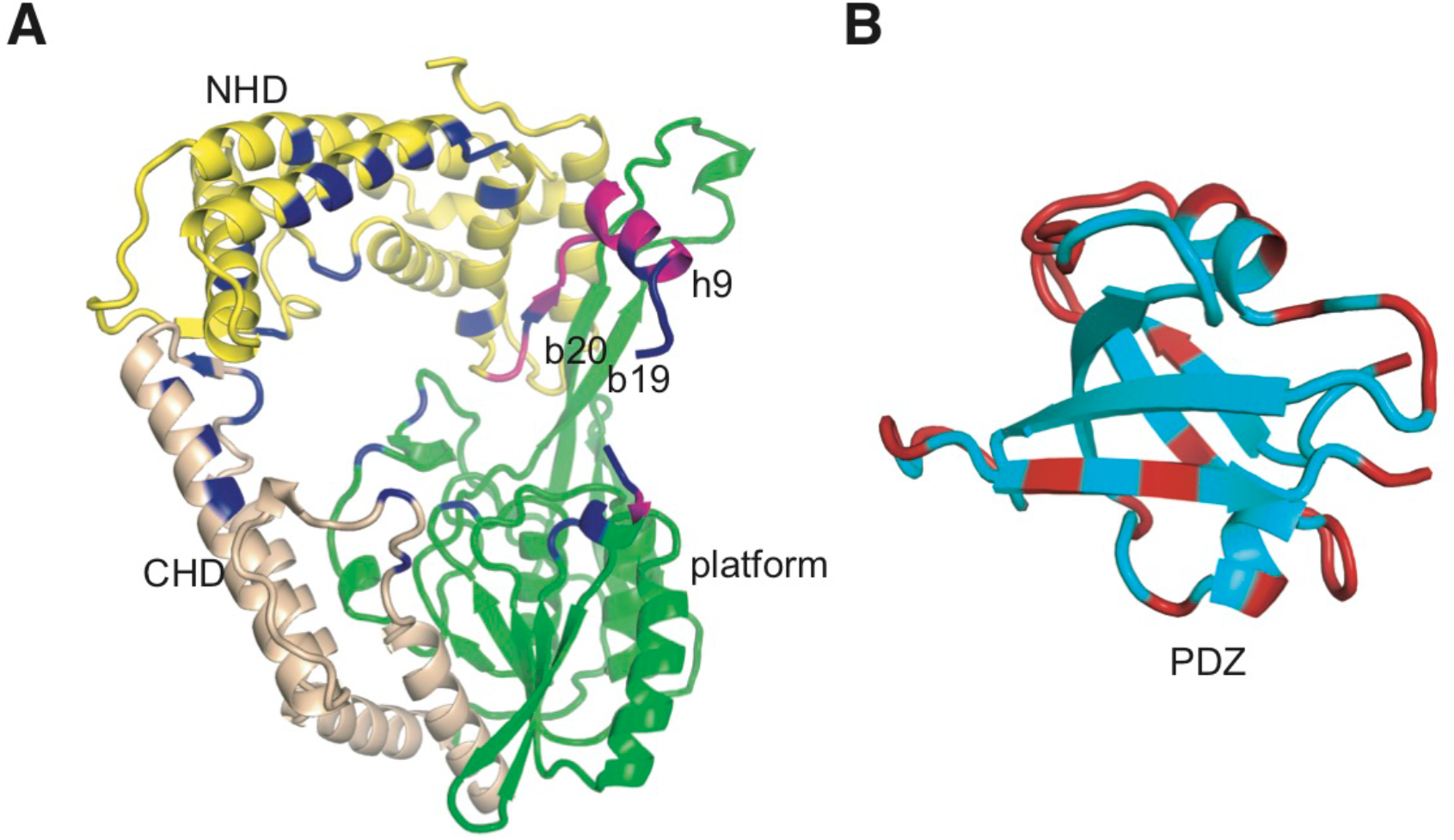
Distribution of the interacting residues of the PDZ domain and the bowl-like scaffold of Prc in the resting state. **(A)** A view of the bowl-like structure of Prc-S452I/L252Y highlighting the residues (in blue) interacting with the PDZ domain (omitted for clarity). **(B)** The residues of the PDZ domain that interact with the bowl-like body of Prc in the resting state are colored in a ribbon model.

**Figure S2.**
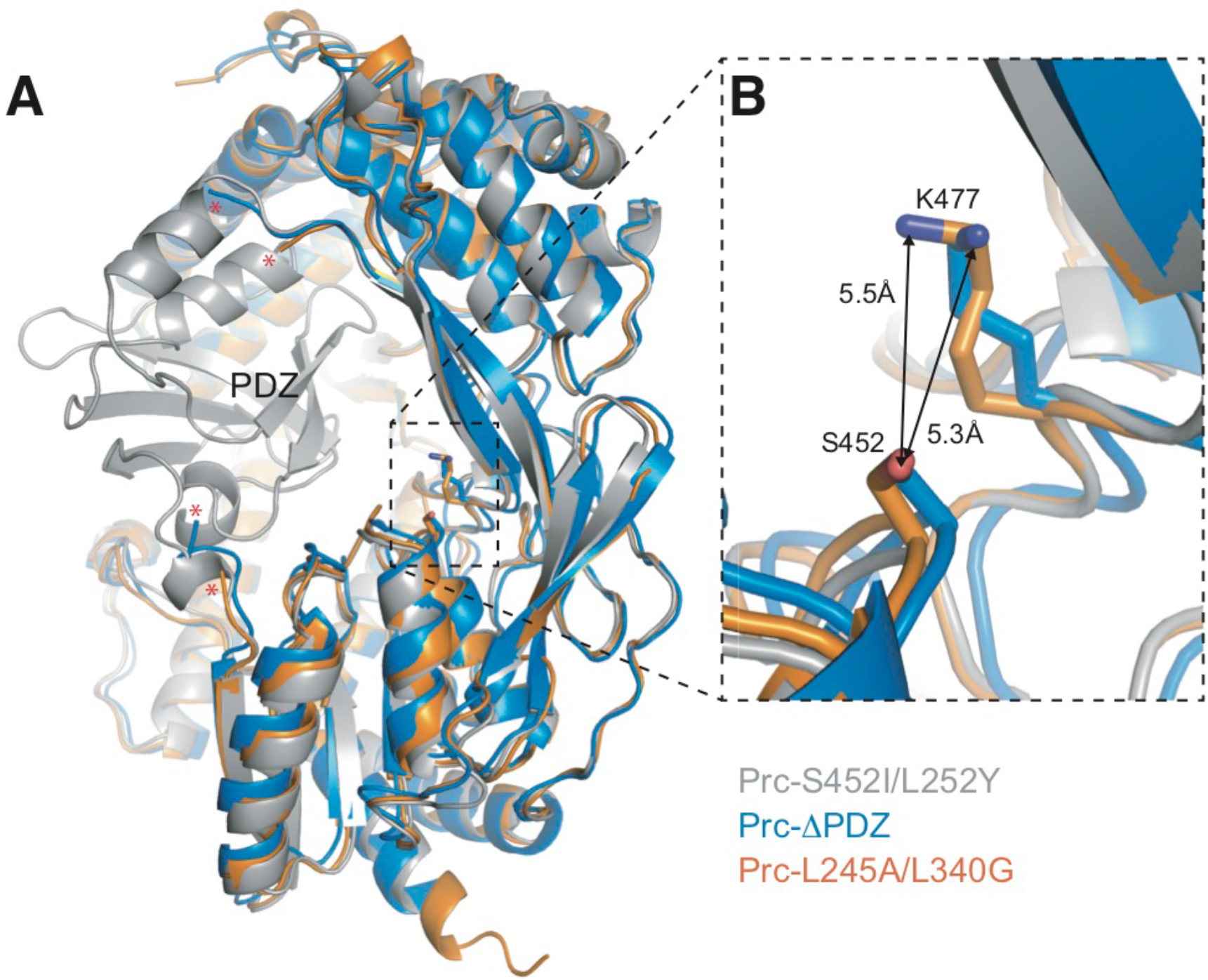
Structures of Prc without the PDZ domain or the hydrophobic sensor. **(A)** Structures of NlpI-Prc-ΔPDZ and NlpI-Prc-L245A/L340G (NlpI is omitted for clarity) are superimposed with Prc-S452I/L252Y. The chain breaks owing to the disordered helix h9 and the PDZ domain, which are missing in the electron-density map, are marked by red asterisks. **(B)** Zoom-in window showing the separated catalytic Lys477 and Ser452 residues.

**Figure S3.**
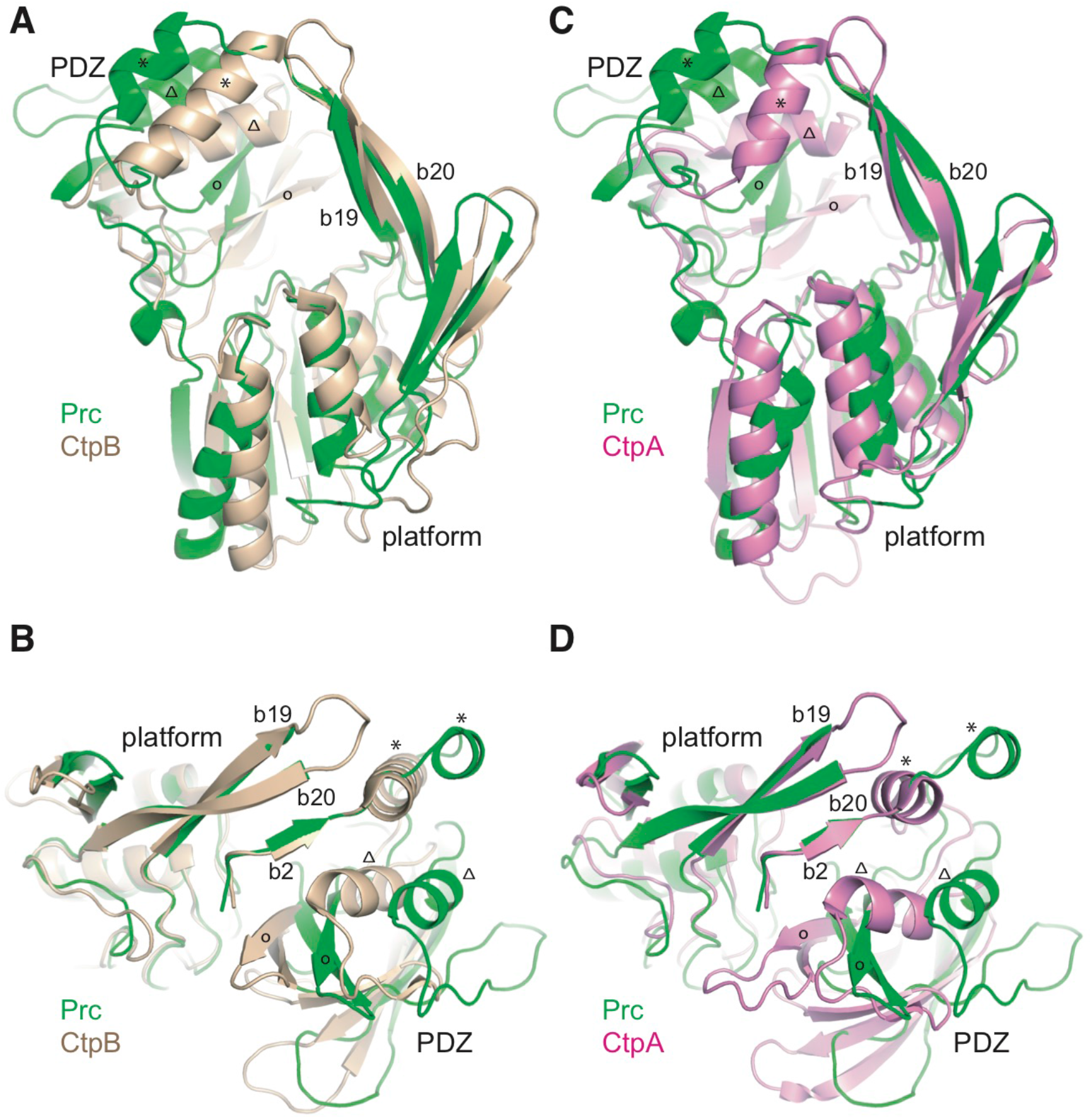
Prc has a larger open gate than the C-terminal processing proteases CtpA and CtpB. **(A)** Superposition of the protease domains of Prc and CtpB in the resting state, aligned based on the vault strand b2 of Prc. The gate helix h9 of Prc and the corresponding helix of CTP proteases are indicated by asterisks. Substrate C-terminal peptide is anchored in between the conserved α-helix and β-strand of the PDZ domain, marked by open triangles and circles, respectively. **(B)** Top view of the aligned Prc and CtpB structures. **(C)** Superposition of the protease domains of Prc and CtpA in the resting state, aligned based on the vault strand b2. **(D)** Top view of the aligned Prc and CtpA.

**Figure S4.**
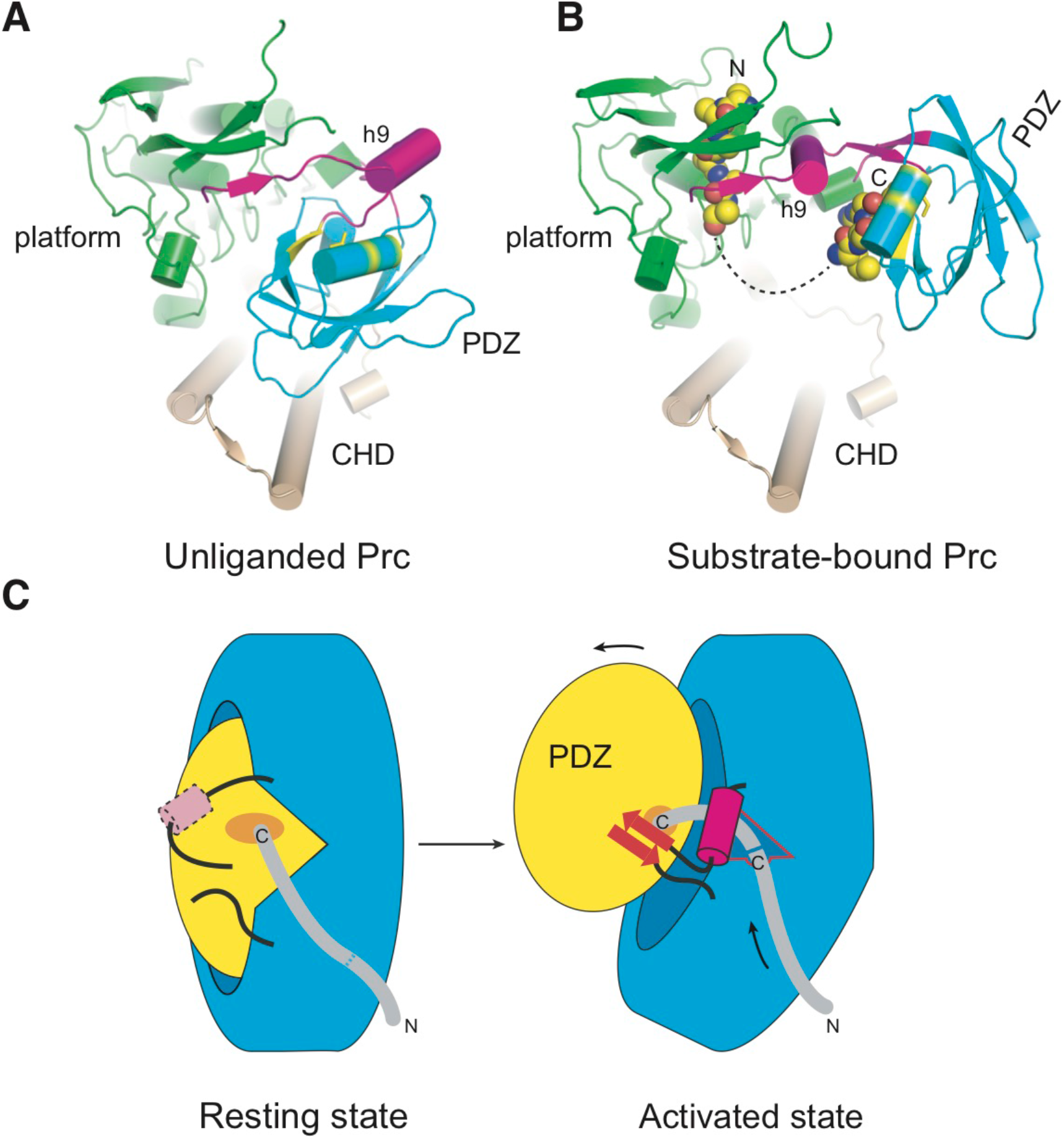
The pulley-like helix h9 in the unliganded resting and the liganded activated states. **(A-B)** Top views of the unliganded resting Prc-S452I/L252Y (A) and substrate-bound activated Prc-K477A (B). The NHD domain has been removed for clarity. The coloring scheme is the same as in Fig. 2. The bound polypeptide chains are shown in spheres. The dashed curve depicts a plausible connection between the cleavage-site (left) and C-terminal (right) peptides bound to Prc-K477A. **(C)** Cartoons illustrating the disorder-to-order transition of helix h9 and its pulley-like role associated with *in cis* substrate-triggered mechanical operation of Prc. The dashed line on the grey-colored substrate C-terminal tail denotes a potential cleavage site (left), which is translocated to the active site and cleaved (denoted by the solid blue line) by the coordinated movements of helix h9 and the liganded PDZ domain in the activated state (right).

### Supplemental Movie

**Movie S1. Ligand-triggered motion and activation of Prc. Related to Figure 4.**

Movie with a sequence showing the overall structure of unliganded Prc, its association with NlpI, and the conformational change, induced by binding of the substrate C-terminus to the PDZ domain, from the unliganded resting state (this work; PDB code 6IQR_a) to the substrate-bound activated state (PDB code 5WQL_c). Also shown in the sequence is a modeled substrate C-terminal peptide (LSRS), derived from the substrate MepS (Su et al., 2017). Helix h9 is colored in magenta. The PDZ domain is colored in yellow. The substrate sensor (Leu340) and the catalytic dyad (Lys477 and Ser452) residues are shown in spheres. Movie and morphing were created with UCSF Chimera (Pettersen et al., 2004). The morphing speed was chosen arbitrarily for illustrative purpose.

**Table S1.** Data collection and Refinement Statistics.

|  | Prc-S452I/L252Y | NlpI-Prc-S452I/L252Y |
| --- | --- | --- |
| Space group | P3<sub>2</sub>21 | P2<sub>1</sub>2<sub>1</sub>2<sub>1</sub> |
| Cell dimensions |  |  |
| a, b, c (Å) | 129.28, 129.28, 236.59 | 118.94, 143.88, 153.59 |
| α, β, γ (°) | 90, 90, 120 | 90, 90, 90 |
| **Data collection** |  |  |
| Wavelength (Å) | 1.00000 | 0.99984 |
| Resolution (Å) | 50~3.42 (3.54~3.42) | 30~2.80 (2.90~2.80) |
| Total observations | 136,067 | 509,147 |
| Unique reflections | 31,494 (3,117) | 64,129 (6,401) |
| R<sub>means</sub> (%) | 14.5 (94.2) | 15.4 (74.8) |
| R<sub>merge</sub> (%) | 7.5 (65.1) | 10.2 (61.5) |
| R<sub>pim</sub><sup>b</sup> (%) | 6.9 (44.6) | 5.5 (26.3) |
| Completeness (%) | 99.4 (100) | 98.3 (99.4) |
| I/σ (I) | 13.8 (2.1) | 13.9 (3.30) |
| Redundancy | 4.3 (4.5) | 7.9 (6.9) |
| **Refinement** |  |  |
| Resolution (Å) | 29.97~3.42 (3.50~3.42) | 29.98~2.80 (2.87~2.80) |
| R<sub>work</sub>/R<sub>free</sub> (%) | 23.9/28.7 | 22.2/27.8 |
| No. of Reflections | 27,244 | 60,219 |
| No. atoms /B factor (Å<sup>2</sup>) |  |  |
| Protein | 101,143/74.06 | 14,407/56.12 |
| Water | 2/50.00 | 95/39.27 |
| Calcium ion |  | 2/53.09 |
| Rms deviations |  |  |
| Bond lengths (Å) | 0.014 | 0.012 |
| Bond angles (°) | 1.95 | 1.88 |
| Ramachandran statistics (%) |  |  |
| most favored | 89.7 | 91.6 |
| additionally allowed | 9.0 | 8.1 |
| generously allowed | 0.9 | 0.3 |
| disallowed regions | 0.4 | 0.0 |
| disordered regions | chain B 533-541 | chain C 242-243<br>chain D 242-243, 324 |
| PDB code | 6IQR | 6IQQ |

**Table S1 (continued). Data collection and Refinement Statistics**
|  | NlpI-Prc-L245A/L340G | NlpI-Prc-ΔPDZ |
| --- | --- | --- |
| Space group | P2<sub>1</sub>2<sub>1</sub>2<sub>1</sub> | C2<sub>1</sub>2<sub>1</sub>2 |
| Cell dimensions |  |  |
| a, b, c (Å) | 119.34, 144.41, 152.50 | 105.83, 151.08, 148.38 |
| α, β, γ (°) | 90, 90, 90 | 90, 90, 90 |
| **Data collection** |  |  |
| Wavelength (Å) | 1.10000 | 0.99984 |
| Resolution (Å) | 30~2.70 (2.80~2.70) | 50~2.90 (3.00~2.90) |
| Total observations | 351,114 | 111,108 |
| Unique reflections | 69,840 (6,187) | 25,531 (1,987) |
| R<sub>means</sub> (%) | 9.8 (81.3) | 8.8 (27.6) |
| R<sub>merge</sub> (%) | 6.4 (54.8) | 5.7 (20.1) |
| R<sub>pim</sub> (%) | 4.2 (39.4) | 4.2 (14.1) |
| Completeness (%) | 94.7 (85.1) | 95.9 (76.0) |
| I/σ (I) | 13.4 (3.68) | 18.9 (4.4) |
| Redundancy | 5.0 (3.5) | 4.4 (3.4) |
| **Refinement** |  |  |
| Resolution (Å) | 29.91~2.69 (2.76~2.69) | 38.80~2.90 (2.98~2.90) |
| R<sub>work</sub>/R<sub>free</sub> (%) | 20.5/24.5 | 19.2/23.3 |
| No. of Reflections | 65,488 | 22,490 |
| No. atoms /B factor (Å<sup>2</sup>) |  |  |
| Protein | 12,841/54.70 | 6434/53.81 |
| Water | 51/39.53 | 11/42.17 |
| Rms deviations |  |  |
| Bond lengths (Å) | 0.021 | 0.015 |
| Bond angles (°) | 2.05 | 2.08 |
| Ramachandran statistics (%) |  |  |
| most favored | 92.0 | 89.3 |
| additionally allowed | 7.6 | 10.4 |
| generously allowed | 0.1 | 0.3 |
| disallowed regions | 0.3 | 0.0 |
| disordered regions | chain C 233-342<br>chain D 233-342 | Chain B 235-247 |
| PDB code | 6IQS | 6IQU |

<sup>a</sup>Highest resolution shell is shown in parenthesis<sup>b</sup>R<sub>pim</sub> is the precision-indicating merging R, which describes the accuracy of the averaged measurement (Weiss,

**Table S2.** Alignment of the resting state structures of Prc.

| Crystal structure | Chain | Two-chain r.m.s.d (Å)* | Cα atoms used for superposition |
| --- | --- | --- | --- |
| NlpI-Prc-S452I/L252Y | C | 0.92/1.16 | 587/637 |
| NlpI-Prc-S452I/L252Y | D | 0.82/1.08 | 591/636 |
| Prc-S452I/L252Y | B | 0.38/0.68 | 547/631 |
| NlpI-Prc-L245A/L340G | C | 0.60/1.51 | 480/529 |
| NlpI-Prc-L245A/L340G | D | 0.66/1.54 | 485/529 |
| NlpI-Prc-ΔPDZ | B | 1.04/1.88 | 519/534 |
| NlpI-Prc-S452I | C | 0.60/0.96 | 503/580 |
| NlpI-Prc-S452I | D | 0.56/1.78 | 491/564 |

\*Backbone r.m.s.d (Å) was calculated over the indicated chain of each crystal structure and chain A in the structure of Prc-S452I/L252Y with and without outlier rejection (5 cycles, a cut-off of 2.0)

**Table S3.**
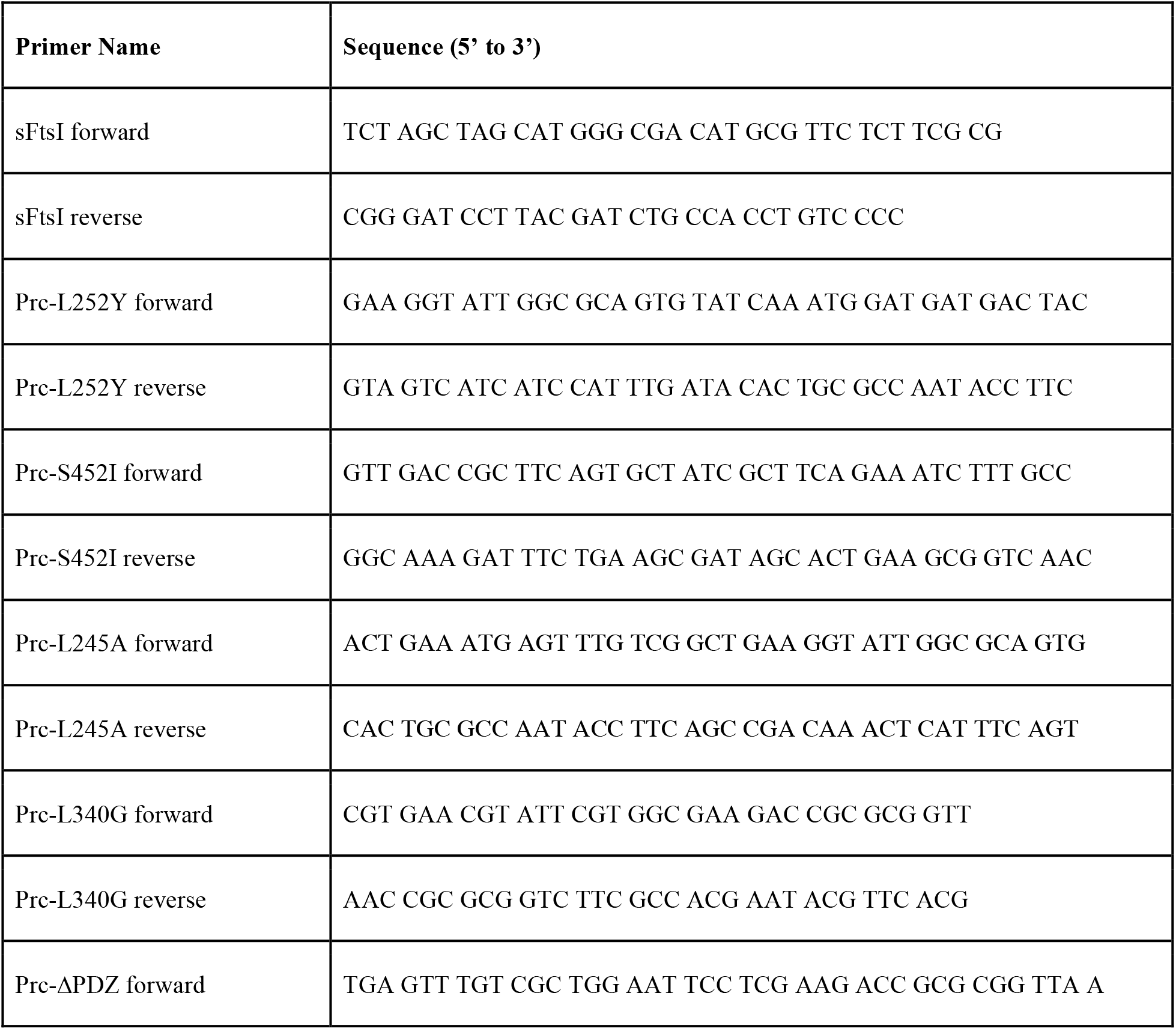
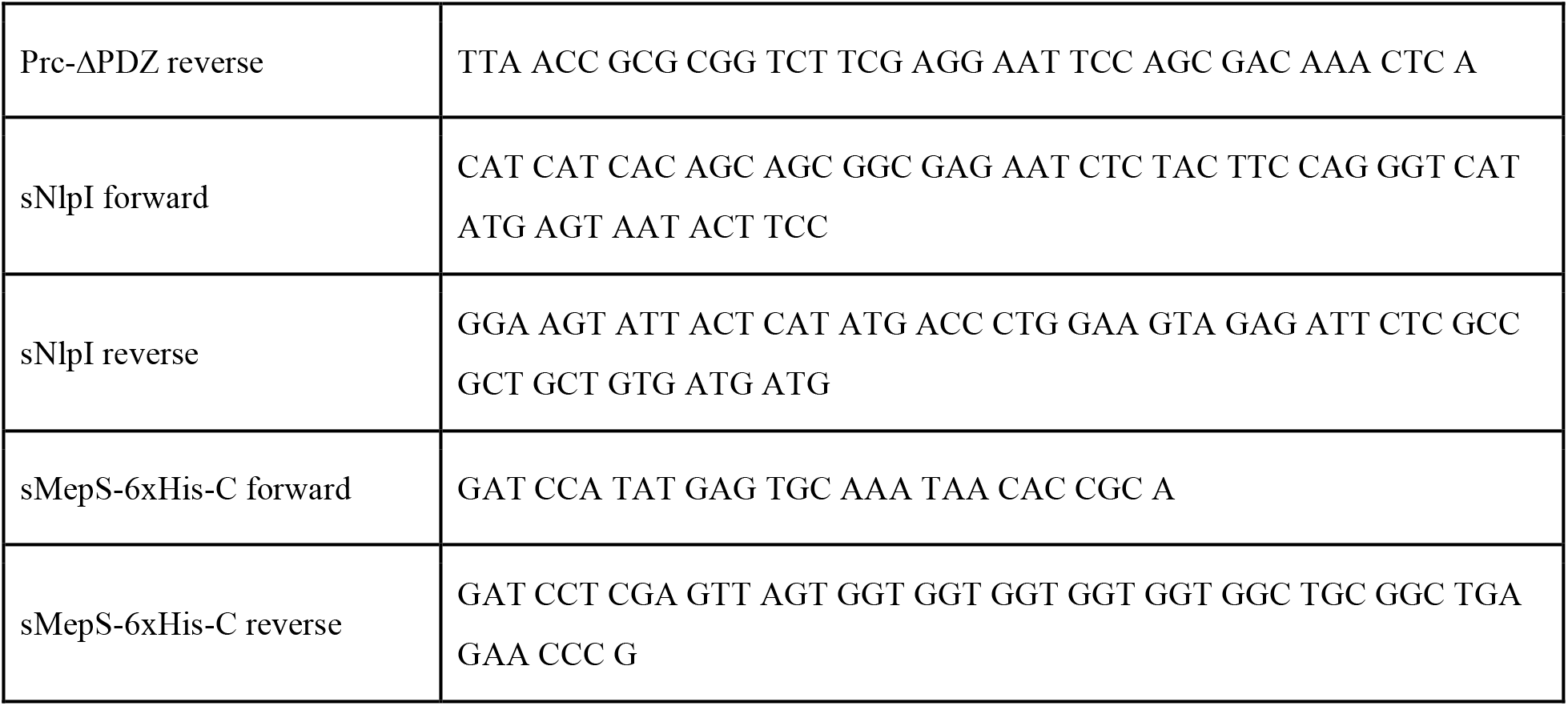
List of primer sequences used in the study.

